# Zika virus NS3 protease and its cellular substrates

**DOI:** 10.1101/2020.09.18.303867

**Authors:** Agnieszka Dabrowska, Aleksandra Milewska, Joanna Ner-Kluza, Piotr Suder, Krzysztof Pyrc

## Abstract

Zika virus is a flavivirus discovered in 1947, but the association between Zika virus infection and brain disorders was not demonstrated until 2015 in Brazil. Infection mostly poses a threat to women during pregnancy, since it may cause microcephaly and other neurological dysfunctions in the developing fetus. However, infection is also associated with Guillain-Barré syndrome. The nonstructural NS3 protein is essential for virus replication because it helps to remodel the cellular microenvironment. Several reports show that this protease can process cellular substrates and thereby modify cellular pathways that are important for the virus. Herein, we explored some of the targets of NS3, but we could not confirm the biological relevance of its protease activity. Thus, although mass spectrometry is highly sensitive and useful in many instances, being also able to show directions, where cell/virus interaction occurs, we believe that biological validation of the observed results is essential.

## Introduction

Zika virus is a flavivirus discovered in 1947 in primates inhabiting the African Zika forest (1). Although the virus was found to infect humans, for decades it was not considered to be a medical threat due to limited distribution and very mild symptoms associated with infection. However, more than 10 years ago interest in Zika virus began to increase as it became clear that the virus has broadened its geographic distribution, and the first outbreak was reported in the Federated States of Micronesia (2). In 2015, case definition became more precise, and some data suggested that the infection may be more dangerous than previously thought. While the symptoms are relatively mild and include fever, rash, headache, and muscle pain, infection may cause severe sequelae, and it is associated with Guillain-Barré syndrome (3). The infection is most severe in pregnant women, since it interferes with development of the neurological system of the fetus, predominantly resulting in microcephaly. A number of *in vitro* and *in vivo* studies have confirmed these observations (4–10).

Flaviviruses are small, enveloped viruses with positive-strand RNA genome, which is delivered to the target cell as a single-stranded RNA molecule containing a single open reading frame (ORF). This ORF is translated into an immature polyprotein, which is co- and post-translationally cleaved by viral and cellular proteases to yield 10 mature viral proteins; capsid (C), membrane (prM/M), and envelope (E) structural proteins; and seven nonstructural proteins (NS1, NS2A, NS2B, NS3, NS4A, NS4B, and NS5)(11). Cleavage sites processed by the viral serine NS3 protease are located between NS2A/NS2B, NS2B/NS3, NS3/NS4A, and NS4B/NS5. Furin or similar cellular proteases process the prM/M site, while other host cell proteases reportedly cleave C/prM, prM/E, E/NS1, NS1/NS2A, and NS4A/NS4B sites(12). The NS3 protein consists of an N-terminal serine protease domain (~180 amino acids) and a C-terminal region harboring RNA helicase, nucleoside triphosphatase, and 5’ RNA triphosphatase activities. The active site of the N-terminal protease contains a His-Asp-Ser catalytic triad. Furthermore, the small nonstructural NS2B protein serves as an NS3 protease cofactor and anchors it to the endoplasmic reticulum (ER) membrane (13–17). The presence or absence of NS2B affects the tertiary structure, activity, and stability of NS3 (18–20).

Flaviviral proteases are essential for viral replication, hence they are considered promising targets for antiviral agents. Indeed, the development of HCV NS3/NS4A protease inhibitors proved a breakthrough in hepatitis C therapy, and these drugs received U.S. Food & Drug Administration (FDA) and European Medicines Agency (EMA) approval for use in humans (21–23).

Interestingly, flaviviral proteases have also been reported to modify the cellular microenvironment. Cleavage of host proteins may be beneficial for the virus by diminishing the cellular responses, remodeling cellular metabolism, and other mechanisms. Such a strategy is common for viruses, as exemplified by human rhinovirus (HRV) that modulates apoptosis by cleaving receptor-interacting protein kinase-1 (RIPK1) at the noncanonical site, and blocking caspase 8-mediated activation of the pathway (24). Interestingly, the picornaviral protease also processes translation initiation factor eIF4G, part of the cellular translation initiation complex. Targeting of this molecule results in decreased production of cellular proteins but does not affect the production of viral proteins, as picornaviruses use internal ribosome entry sites (IRES) for cap-independent translation. In this way, the viral protease hijacks the cellular protein production machinery (25–27). The NS3/NS4A protease of hepatitis C virus cleaves mitochondrial antiviral signaling protein (MAVS) and TIR domain-containing adapter-inducing interferon-β protein (TRIF) to evade the host cell antiviral response (28–30).

Various targets of the Zika virus protease have been identified. FAM134B (family with sequence similarity 134), ATG16L1 (autophagy-related protein 16-1), eIF4G1 (eukaryotic translation initiation factor 4 gamma 1), and Septin-2 are among the most interesting targets. Except for FAM134B, these targets were identified using mass spectrometry methods, which provide unrivalled sensitivity and are capable of cell proteome studies. However, careful analysis of the reported data showed that none of the protein targets are shared between the published studies. Thus, we explored the reported data in detail by employing classical approaches such as western blotting, as well as functional approaches based on the activity of particular pathways. Herein, we present an example of such a study, in which we failed to confirm protease-mediated or virus-related degradation of the eIF4G1 protein. We were also unable to confirm the beneficial effects of decreasing eIF4G1 on viral replication.

## Materials and Methods

### Cell culture

293T cells (ATCC CRL-3216; human embryonic kidney cells), A549 cells (ATCC CCL-185; lung carcinoma cells), Vero cells (ATCC CCL-81; African green monkey kidney cells), and U251 cells (human glioblastoma cell line) were maintained in Dulbecco’s modified Eagle’s medium (DMEM; Corning, Poland) supplemented with 3% fetal bovine serum (FBS; heat-inactivated; Thermo Scientific, Poland), 100 μg/ml streptomycin, 100 U/ml penicillin (Sigma-Aldrich, Poland), and 5 μg/ml ciprofloxacin. Cells were maintained at 37°C under 5% CO_2_.

### Virus strains, preparation, and titration

ZIKV H/PF/2013 (acquired from European Virus Archive), ZIKV H/PAN/2016 (BEI resources), ZIKV R116265 Human 2016 Mexico (BEI resources), ZIKV Mosquito Mex 2-81 (BEI resources), ZIKV PRVABC59 (BEI resources), ZIKV MR766 (BEI resources), ZIKV IB H 30656 (BEI resources), ZIKV FLR (BEI resources), ZIKV R103451 Human 2015 Honduras (BEI resources), ZIKV P 6-740 Malaysia 1966 (BEI resources), and ZIKV DAKAR 41524 (BEI resources) strains were employed in this work.

Virus stocks were generated by infection of Vero cells. At 3 days post-infection (p.i.) at 37°C, virus-containing medium was collected and titrated. As a control, mock-infected Vero cells were subjected to the same procedure. Virus and mock aliquots were stored at −80°C. Virus titration was performed on confluent Vero cells in a 96-well plate according to the method described by Reed-Muench(31). Briefly, cells infected with serially diluted virus were incubated at 37°C for 3 days, and the occurrence of a cytopathic effect (CPE) was monitored.

### Plasmids

The region encoding the NS2B-NS3^WT^ protein was amplified by PCR using a cDNA template generated from H/FP/2013 Zika virus and appropriate primers (5’ ATG CGG TAC CGC CAC CAT GGG CAG CTG GCC CCC TAG CGA A 3’; 5’ AGC CGG TAC CCT ATC TTT TCC CAG CGG CAA ACT CC 3’). The resulting product was digested with *Not*I-HF and *Kpn*I-HF (New England Biolabs), gel-purified, and cloned into the pBudCE4.1 vector (pBudCE4.1_NS3^WT^). The plasmid encoding the inactive NS2B-NS3^S135A^ protease (pBudCE4.1-NS3^S135A^) was obtained using the pBudCE4.1_NS3^WT^ template by employing the QuickChange PCR technique with appropriate primers (5’ GGA ACT GCC GGA TCT CCA ATC CTA GAC AAG 3’; 5’ AGA TCC GGC AGT TCC TGC TGG GTA ATC CAG 3’) to change the serine residue at amino acid (aa) position 135 to alanine. The obtained plasmids were verified by DNA sequencing.

### Plasmid transfection

293T cells were maintained as described above. Cells were seeded in 6- or 24-wells plates (TPP, Switzerland) and cultured for 24 h. When 60% confluency was reached, cells were transfected using polyethyleneimine (PEI; Sigma-Aldrich, Poland). For transfection in 6-wells plates, 4 μg plasmid DNA was mixed with 250 μl Opti-MEM medium (Thermo Scientific) and 4 μg PEI. For transfection in 24-well plates, 1 μg/well plasmid DNA was mixed with 100 μl Opti-MEM medium with 1 μg PEI. After a 30 min incubation at room temperature, the mixture was added dropwise onto cells. Four hours later, the supernatant was discarded, fresh medium was added, and cells were further incubated at 37°C.

A549 cells were maintained as described above. Cells were seeded in 6-wells plates and cultured for 24 h. When 80% confluency was reached, cells were transfected with Lipofectamine 2000 (Thermo Scientific) according to the manufacturer’s protocol. Briefly, 2.5 μg plasmid DNA was mixed with 300 μl Opti-MEM medium with 5 μl Lipofectamine 2000. After a 5 min incubation at room temperature, the mixture was added dropwise onto cells. Four hours later, the supernatant was discarded, fresh medium was added, and cells were further incubated at 37°C.

For expression of active and inactive virus protease, pBudCE4.1-NS3^WT^ or pBudCE4.1-NS3^S135A^ plasmids were employed, respectively. For eIF4G1 overexpression, cells were transfected with pcDNA3 HA eIF4GI plasmid or control green fluorescent protein (GFP)-expressing plasmid (pMAX-GFP plasmid, Lonza). pcDNA3 HA eIF4GI (1–1599) was a gift from Nahum Sonenberg (Addgene plasmid #45640; http://n2t.net/addgene:45640; RRID Addgene_45640). The efficiency of expression was verified by western blotting.

### siRNA transfection

For small interfering RNA (siRNA) transfection, A549 cells were maintained as described above. Cells were seeded in 24-wells plates, and siRNA was transfected once the confluency reached 80% using RNAiMAX Lipofectamine (Thermo Scientific), according to the manufacturer’s protocol. Next, 5 pmol eIF4G1 siRNA (Sigma-Aldrich; Cat. No EHU066831) or control scrambled RNA (Santa-Cruz Biotechnology; Cat. No sc-44237) was mixed in 125 μl Opti-MEM medium containing 3.5 μl transfection reagent. After a 5 min incubation at room temperature, the mixture was added to cells dropwise. The efficiency of eIF4G1 silencing was verified at 24–72 h post-transfection using western blotting.

### Virus infection

293T cells, A549 cells, and U251 cells were seeded in 6-wells plates and cultured at 37°C. When 90–100% confluency was reached, cells were inoculated with ZIKV at 2000 TCID_50_/ml for 293T cells or 400 TCID_50_/ml for U251 and A549 cells. Mock cultures were inoculated with an identical volume of mock samples. All cultures were incubated for 2 h at 37°C under 5% CO_2_ in DMEM medium supplemented with 2% FBS, 100 μg/ml streptomycin, and 100 IU/ml penicillin. After incubation, cells were washed twice with phosphate-buffered saline (PBS) and incubated as described above. At 3 days p.i., culture supernatants were collected, viral RNA was isolated, and the yield was quantified by reverse transcription quantitative PCR (RT-qPCR). Also, cells were collected for western blotting analysis.

### SUnSET-puromycin assay

A549 and U251 cells were maintained as described above. Cells were seeded in 12-well plates and cultured for 48 h. When 80% confluency was reached, cells were inoculated with the Mexico ZIKV strain at 400 TCID_50_/ml (or an identical volume of the mock culture). Alternatively, 10 μg/ml or 5 μg/ml of the translation inhibitor cycloheximide (CHX; stock solution 100 mg/ml; Sigma-Aldrich) was added to A549 cells and U251 cells, respectively, or 10 μM eIF4G1 inhibitor (4EGI-1; stock solution 10 mM; Biotechne)(32) was added. At 48 h p.i., cells were washed twice with PBS and incubated in unsupplemented DMEM for 2 h at 37°C. The supernatant was discarded, fresh DMEM medium supplemented with 3% FBS, and 1 μM puromycin (stock solution; Merck) was added, and cells were further incubated for 30 min at 37°C. Subsequently, cells were collected for western blotting analysis.

### SDS-PAGE and western blotting

Cells grown in 6-well, 12-well, or 24-well plates were lysed for 30 min on ice in 200 μl, 100 μl, or 50 μl RIPA buffer (50 mM TRIS, 150 mM NaCl, 1% Nonidet P-40, 0.5% sodium deoxycholate, 0.1% SDS, pH 7.5), respectively. Subsequently, samples were centrifuged (10 min at 13,000 × g, 4°C), and the pelleted cell debris was discarded. The total protein concentration in each sample supernatant was quantified using the bicinchoninic acid (BCA) method (Pierce BCA Protein Assay Kit; Thermo Scientific) according to the manufacturer’s protocol. Supernatants were mixed 5:1 with denaturing buffer (202.5 mM TRIS pH 6.8, 10% SDS, 15% β-mercaptoethanol, 30% glycerol, 0.3% Bromophenol Blue) and boiled at 95°C for 5 min.

For detection of proteins, lysates were loaded and separated on 12% polyacrylamide gels, and separated by SDS-PAGE over 2 h at 120 V. BlueStar Plus Prestained Protein Markers (NIPPON Genetics, Germany) were used for reference. Subsequently, gels were subjected to wet electrotransfer onto methanol-activated polyvinylidene difluoride (PVDF; GE Healthcare, Poland) membranes in 25 mM TRIS, 192 mM glycine, 20% methanol buffer for 1 h at 100 V. Following transfer, nonspecific binding sites were blocked with 5% skimmed milk (BioShop, Canada) in TRIS-buffered saline (20 mM TRIS, 0.5 M NaCl, pH 7.5) supplemented with 0.05% Tween 20 (TBS-T) by overnight incubation at 4°C. To detect NS3 protein, membranes were incubated with a rabbit anti-NS3 antibody (1:1000, GeneTex, USA) followed by a secondary goat anti-rabbit antibody (1:20000; Dako, Denmark) conjugated with horseradish peroxidase (HRP). To detect the eIF4G1 protein, membranes were incubated with a rabbit anti-eIF4G1 antibody (1:1000; Thermo Scientific) followed by a secondary goat anti-rabbit antibody (1:20000; Dako) conjugated with HRP. For puromycin detection, membranes were incubated with mouse anti-puromycin antibody (1:10000; Merck) followed by a secondary rabbit antimouse antibody (1:20000; Dako) conjugated with HRP. To detect GAPDH protein, membranes were incubated with a rabbit anti-GAPDH antibody (1:5000; Cell Signaling) followed by a secondary goat anti-rabbit antibody (1:20000; Dako) conjugated with HRP. All antibodies were diluted in 1.5% skimmed milk in TBS-T. The signal was developed using Immobilon Western Chemiluminescent HRP Substrate (Millipore, Poland) and recorded with a ChemiDoc Imaging System (Bio-Rad, Poland).

### Isolation of nucleic acid and reverse transcription

Viral RNA was isolated from 100 μl cell culture supernatant using a viral DNA/RNA Isolation Kit (A&A Biotechnology, Poland) according to the manufacturer’s protocol. Reverse transcription was carried out using a High Capacity cDNA Reverse Transcription Kit (Thermo Scientific) according to the manufacturer’s protocol. cDNA samples were prepared in 10 μl volumes using a High Capacity cDNA Reverse Transcription Kit (Thermo Scientific) according to the manufacturer’s instructions. The reaction was carried out for 10 min at 25°C, 120 min at 37°C, and 5 min at 85°C.

### Quantitative PCR (qPCR)

Zika virus RNA yields were assessed using real-time PCR on a 7500 Fast Real-Time PCR instrument (Thermo Scientific, Poland). ZIKV cDNA was amplified in a reaction mixture containing 1× TaqMan Universal PCR Master Mix (RT-PCR mix; A&A) in the presence of FAM/TAMRA (6-carboxyfuorescein/6-carboxytetramethylrhodamine) probe (5’ CGG CAT ACA GCA TCA GGT GCA TAG GAG 3’; 100 nM) and primers (5’ TTG GTC ATG ATA CTG CTG ATT GC 3’ and 5’ CCT TCC ACA AAG TCC CTA TTG C 3’; 450 nM each). The reaction was carried out for 2 min at 50°C and 10 min at 92°C, followed by 40 cycles at 92°C for 15 s and 60°C for 1 min. DNA standards were subjected to qPCR along with the cDNA. Rox was used as a reference dye.

## Results

### Expression of Zika virus protease in eukaryotic cells

In this study, we expressed and purified full-length NS2B-NS3 protein without linkers. The NS2B-NS3 protein and its inactive Ser135Ala variant were expressed from pBudCE4.1 plasmids in 293T (left panel) and A549 (right panel) cells. Western blotting with anti-NS3 antibody (full-length recombinant Zika virus NS3 protein was used as a positive control) detected NS2B-NS3^WT^ and NS2B-NS3^S135A^ in cell lysates prepared from cells collected at 48 h post-transfection. The band corresponding to NS3/NS2B was expected to migrate at ~82 kDa, but the active protease should undergo autocatalytic processing to yield the mature ~68 kDa NS3 protein (**Fig. 1A**). Processing should also result in the generation of the smaller 14 kDa NS2B (**Fig. 1B**). All fragments were observed as expected, confirming the activity of the protease, and protein expression was efficient in both cell lines.

**Fig. 1.**
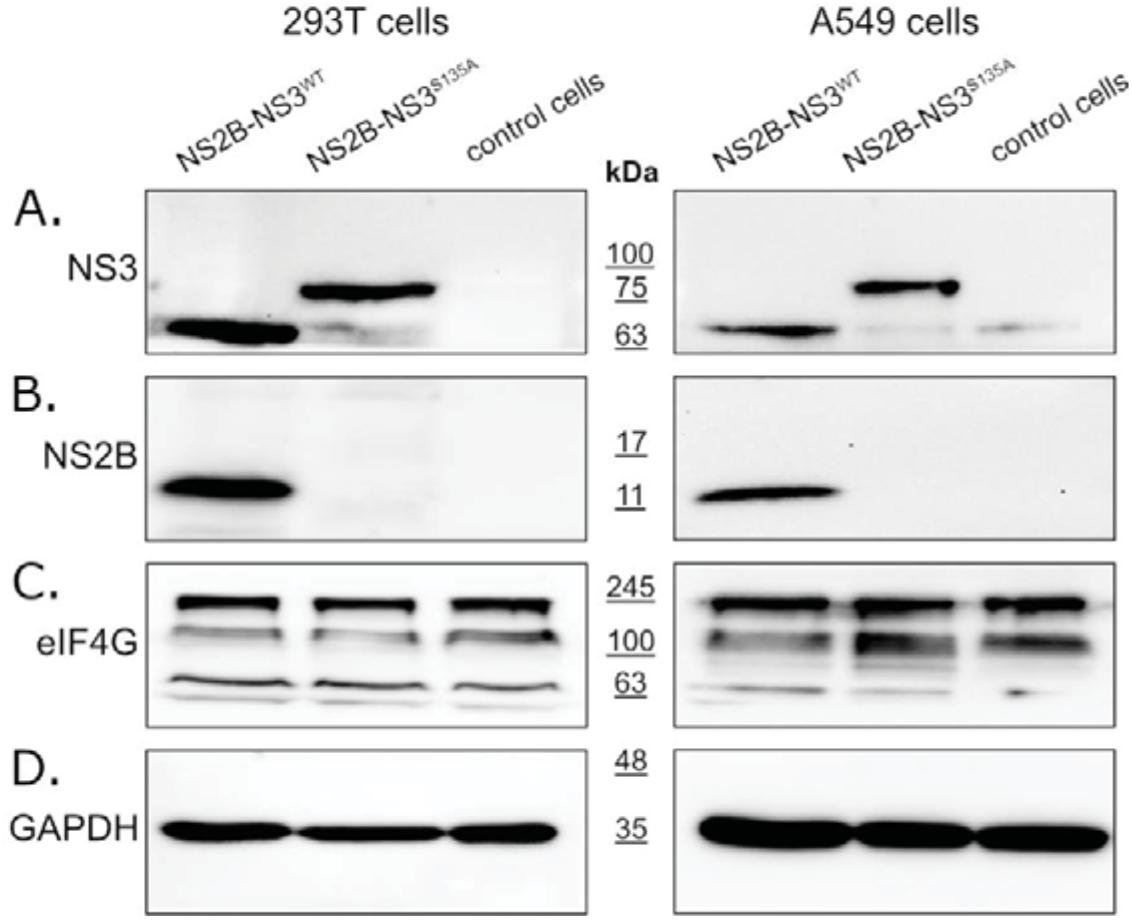
Expression of active and inactive NS2B-NS3 protease from Zika virus in eukaryotic cells does not affect eIF4G1 protein levels. 293T (left panel) and A549 (right) cells expressing NS2B-NS3^WT^ or NS2B-NS3^S135A^ were assessed alongside control cells. Cells were harvested at 48 h after transfection, lysed, and virus lysates were analyzed by western blotting. Virus infection was confirmed by the presence of NS3 (**A**) and NS2B (**B**) proteins. Anti-eIF4G1 antibody was used to detect and compare changes in eIF4G1 protein abundance in cells expressing active or inactive protease, relative to control cells (**C**). The GAPDH protein was used as a reference to ensure that identical amounts of proteins were present in each sample (**D**).

### NS3 protease does not affect eIF4G1 levels

To verify whether the ZIKV NS3 protease has any effect on eIF4G1 levels, cells expressing either active or inactive protease were analyzed used western blotting, alongside control cells lacking the protease. The results (**Fig. 1C**) showed that eIF4G1 migrated at ~188 kDa, and isoforms with lower molecular masses were also visible. While we observed high variability in eIF4G1 content depending on the culture time, temperature, and general cell conditions, there were no differences in protein abundance in cells expressing active or inactive protease, or control cells (data not shown), and this was the case for both 293T and A549 cells.

### Zika virus infection does not result in altered eIF4G1 levels

Since eIF4G1 levels were not affected by the expression of NS3 protease, we assessed whether the eIF4G1 protein is cleaved or degraded during virus infection using ZIKV-infected and mock-infected 293T and A549 cells. First, we confirmed virus replication in cells through NS3 and NS2B protein expression using western blotting **(Fig. 2A, B)**. Levels of the eIF4G1 protein were then assessed in virus-infected and mock-inoculated cells, but there were no differences in eIF4G1 protein abundance.

**Fig. 2.**
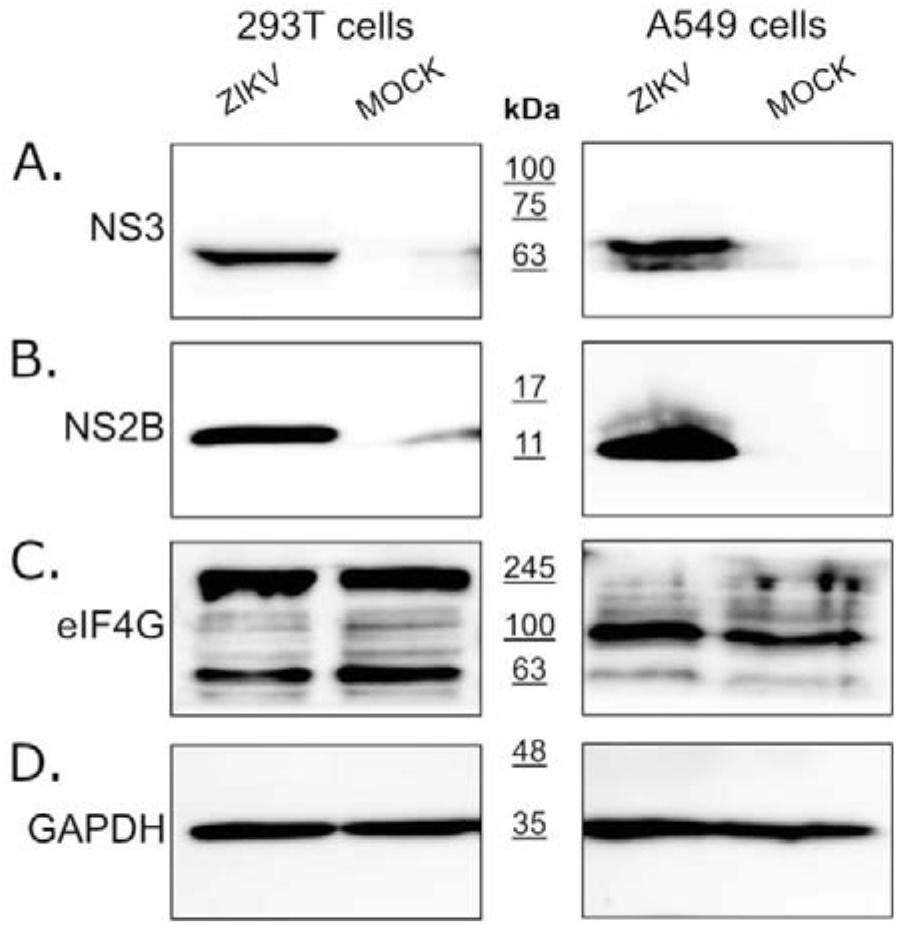
Expression of the eIF4G1 protein in ZIKV-infected and mock-infected cells. 293T (left panel) and A549 (right panel) cells were infected with ZIKV, or inoculated with mock or virus lysates, and analyzed by western blotting. Virus infection was confirmed by the presence of NS3 (**A**) and NS2B (**B**) proteins. The eIF4G1 protein was detected using anti-eIF4G1 antibody (**C**). The GAPDH protein was used as a reference to ensure that identical amounts of proteins were present in each sample (**D**).

### ZIKV does not hamper production of cellular proteins by altering levels of transcription factors

It was suggested that ZIKV NS3 cleaves eIF4G1 to redirect the cellular machinery toward viral protein production, which may be independent of cellular transcription factors(33). To test this hypothesis, the host protein synthesis efficiency was evaluated using surface sensing of translation (SUnSET) assays to measure protein synthesis in cultured cells (34). Puromycin can mimic the aminoacyl end of aminoacyl-tRNAs, and it is partially incorporated in synthesized proteins. The incorporation rate reflects the rate of mRNA translation. Puromycin incorporation was detected by western blotting using anti-puromycin antibodies. Two reference inhibitors were also tested: the protein synthesis inhibitor cycloheximide (CHX) and the eIF4G1-specific inhibitor 4EGI1 **(Fig. 3A)**. In samples treated with either CHX or 4EGI1, the synthesis of proteins was significantly hampered, but protein synthesis in ZIKV-infected cells was not altered. The cytotoxicity of the inhibitors was also evaluated (**Fig. 3B**).

**Fig. 3.**
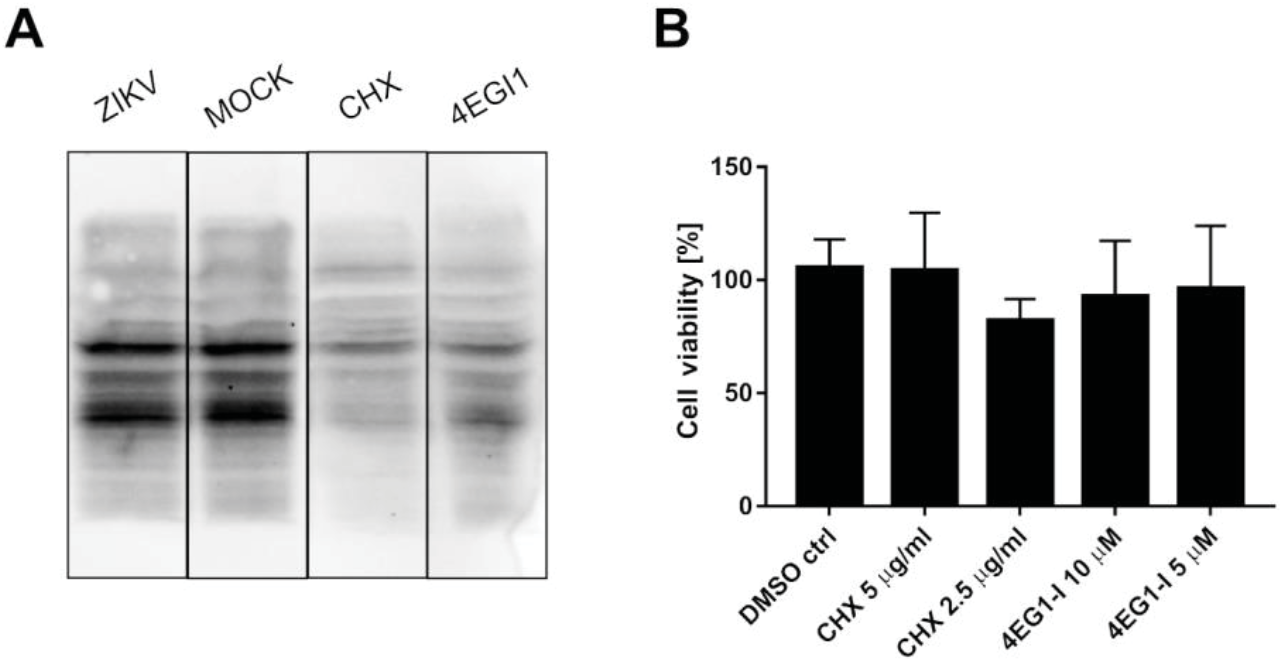
ZIKV infection does not inhibit the translation of host proteins. SUnSet assays were performed on ZIKV- and mock-infected A549 cells, and cells treated with cycloheximide or 4EGI1 inhibitors. The presence of puromycin protein was detected using anti-puromycin antibody (**A**). Cell viability was evaluated relative to control cells treated with DMSO alone. The assay was performed in triplicate, and average values with standard errors are presented (**B**).

### Overexpression of eIF4G1 does not limit ZIKV replication

To verify whether eIF4G1 expression negatively regulates replication of ZIKV, 293T cells were transfected with a plasmid encoding eIF4G1 (or GFP as a control). Protein levels were verified using western blotting **(Fig. 4A)**, and ZIKV replication was evaluated in non-transfected cells, GFP-expressing cells, and eIF4G-expressing cells. Different strains of Zika virus were used to ensure that any effect is not limited to a single lineage. Cells were infected, and at a single timepoint cell culture supernatants were collected for RNA isolation and subsequent RT-qPCR assessment of the virus yield. Although virus yields varied depending on the virus strain, no inhibition of virus replication was observed for any of the tested strains. These results show that NS3-mediated loss of function of the eIF4G1 protein was not beneficial for virus replication **(Fig. 4B)**.

**Fig. 4.**
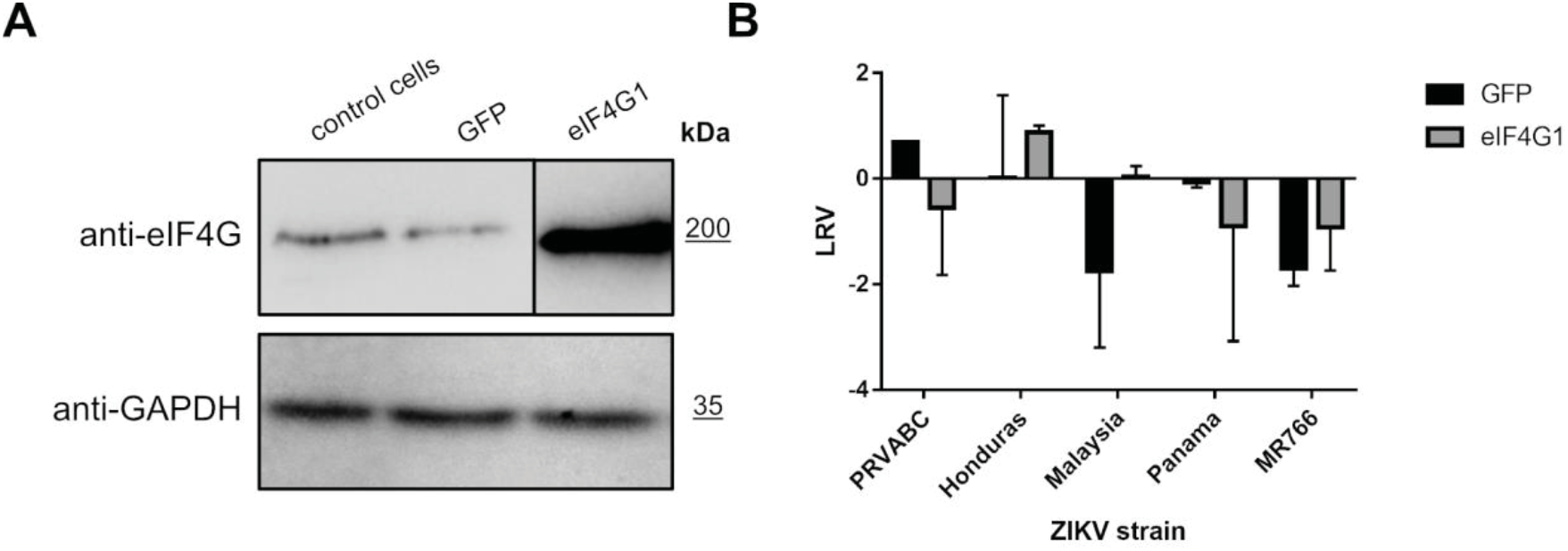
ZIKV replication in eIF4G1-overexpressing cells. Western blotting analysis (anti-eIF4G1 antibodies) was performed on 293T cells transfected with plasmid encoding eIF4G1 or GFP. The GAPDH protein was used as a reference to ensure that identical amounts of proteins were present in each sample (**A**). ZIKV virus replication in 293T cells transfected with plasmid encoding eIF4G1 or GFP. The virus yield was assessed by RT-qPCR. The y-axis represents the log reduction value (LRV) in virus yield in treated samples, and the x-axis corresponds to different ZIKV strains. The assay was performed in triplicate, and average values with standard errors are presented (**B**).

### eIF4G supports replication of ZIKV

To further investigate the role of eIF4G1, we silenced its expression in 293T cells and probed ZIKV replication in these cells. Briefly, cultures were transfected with eIF4G1 siRNA or with scrambled siRNA. Silencing was confirmed by western blotting using antibodies specific to eIF4G1 **(Fig. 5A)**. Cells were infected with ZIKV and incubated for 3 days at 37°C, after which culture supernatants were collected, RNA was isolated and reverse transcribed, and virus replication was evaluated by qPCR. Again, we did not observe an increase in virus production; on the contrary, for some strains, silencing led to inhibition of virus replication relative to control cells or cells transfected with scrambled siRNA **(Fig. 5B)**.

**Fig. 5.**
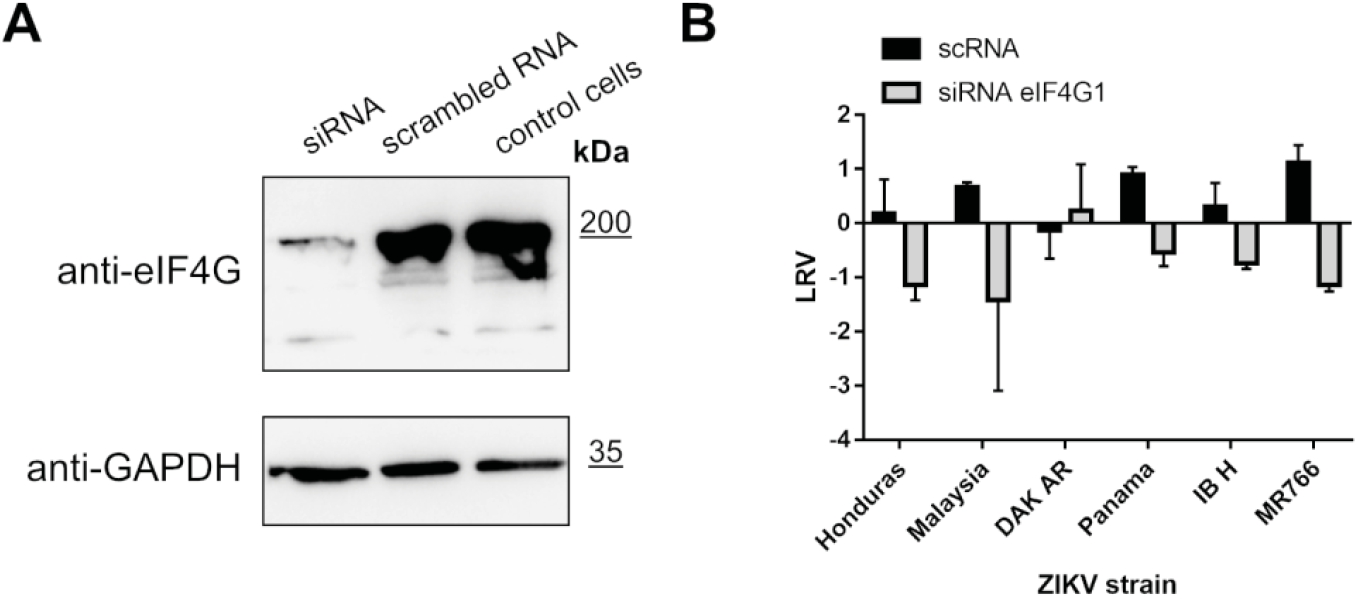
ZIKV replication in cells lacking eIF4G1. A549 cells were transfected with eIF4G1 siRNA or scrambled siRNA. Non-transfected controls were included. The GAPDH protein was used as a reference to ensure that identical amounts of proteins were present in each sample (**A**). ZIKV virus replication in A549 cells transfected with different siRNAs. The virus yield was assessed by RT-qPCR. The y-axis represents the log reduction value (LRV) in virus yield in treated samples, and the x-axis corresponds to different ZIKV strains. The assay was performed in triplicate, and average values with standard errors are presented (**B**).

## Discussion

This study aimed to verify the role of the ZIKV NS3 protein in the remodeling of host cells. NS3 protease is essential for virus replication because it is required for viral protein maturation (35–37). However, viral proteases are generally considered to be highly specific enzymes that co-evolved with the host, and they typically target specific cellular pathways to support viral replication in the host cell, or to block recognition of the virus by the host immune system (38, 39). In the case of ZIKV, some protease targets have been identified, and these are listed in **Table 1**.

**Table 1.** Reported NS3 protease targets in the cell.

|  | PROTEIN | FUNCTION | PROTEASE EXPRESSION | CELLULAR MODEL | IDENTIFICATION | REFERENCES |
| --- | --- | --- | --- | --- | --- | --- |
| 1 | autophagy-related protein 16-1 (ATG16L1) | Autophagy | expression in the prokaryotic system; construct based on 48-100 aa residues of NS2B and 1-178 aa of NS3 (protease domain) | protease substrate verification in the cellular lysate of 293T and A549 cells treated with purified protease; | mass spectrometry; Western blot analysis | (33) |
| 2 | eukaryotic translation initiation factor 4 gamma (eIF4G1) | Translation process | expression in prokaryotic system; construct based on 48-100 aa residues of NS2B and 1-178 aa of NS3 (protease domain) | protease substrate verification in cellular lysate of 293T and A549 cells treated with purified protease; | mass spectrometry; Western blot analysis | (33) |
| 3 | FAM134B | Reticulofagy | expression in eukaryotic cells; the NS2B-NS3 protein coding region was amplified from cDNA produced from HBMEC infected with ZIKV MR766; cells transfection | HBMEC, U2OS, 293T and HeLa cells | Western blots and the fluorescence microscopy analysis | (40) |
| 4 | Septin-2 | Cytokinesis | expression in eukaryotic cells, cells transfection, and transduction with lentiviral vectors; | HeLa and 293T cells and human neural progenitor cells | mass spectrometry; Western blots and fluorescence microscopy analysis, pull-down assay | (41) |
| 5 | disulfide-isomerase A3 (PDIA3) | ER stress response | Expression in eukaryotic cells, cells transfection; construct based on 48-94 aa residues of NS2B and 1-188 aa of NS3 (protease domain) | 293T and A549 cells | mass spectrometry and Western blot analysis | (42) |
| 6 | aldolase A (ALDOA) | glycolysis | Expression in eukaryotic cells, cells transfection; construct based on 48-94 aa residues of NS2B and 1-188 aa of NS3 (protease domain) | 293T and A549 cells | mass spectrometry and Western blot analysis | (42) |
| 7 | Nup98, Nup153 and TPR | Formation of nuclear pore complex | Expression in eukaryotic cells, cells transfection | Huh-7 cells | Western blots and the fluorescence microscopy analysis | (43) |

In the present work, we reviewed the published data and verified these potential protease targets experimentally by measuring changes in the levels of potential cellular targets in the presence of active or inactive protease. Since in some cases the localization or specificity of the protease may differ in the absence of other viral proteins, we also measured the levels of potential NS3 targets in ZIKV-infected cells.

We employed eIF4G1 as a model protease substrate because we believe that processing of this protein has straightforward consequences for both host cells and virus. The eIF4G protein is involved in the translation process by serving as a eukaryotic translation initiation factor. Together with eIF4A and eIF4E, eIF4G forms the EIF4F multi-subunit protein complex, which recognizes the mRNA cap and facilitates the recruitment of mRNA to the ribosome. eIF4G serves mainly as a linker that forms a scaffold for the complex (44). Interestingly, some viruses are reported to target this protein, and thereby rewire the cellular machinery and switch off cellular protein production. For example, coxsackievirus B3 virus-encoded protease cleaves eIF4G1, but the resulting suppression of cellular translation does not affect viral replication, since picornaviruses utilize the IRES rather than cap-dependent translation initiation (25–27). Consequently, the complete protein production machinery serves viral replication. Similarly, for some flaviviruses, it was postulated that the 5’-untranslated region (5’-UTR) may act as an IRES, and NS3 protease encoded in the flaviviral genome may target cellular translation initiation factors (45).

Herein, we first tested whether overexpression of NS2B/NS3 had any effect on levels of the eIF4G1 protein, as reported previously by Hill *et al*. (2018). Notably, the authors of this work performed their analysis using ZIKV protease expressed in a prokaryotic system, which was purified and mixed with cellular lysates from 293T and A549 cells (33). In our current work, we expressed part of the ZIKV genome encompassing the NS2B and NS3 proteins. Our approach allowed us to anchor the NS3 protease in the ER membrane *via* the NS2B cofactor.

Furthermore, using this approach, we were able to monitor whether the protease was active in every experiment because it was autocatalytically (in trans and cis) processing its natural substrate (the NS2B/NS3 junction). The catalytically inactive mutant was used as a negative control. Protein content analysis did not reveal any significant decrease in eIF4G1 protein levels. However, the experimental setup used in the present study may not be entirely appropriate, since it may not accurately recapitulate protease activity and localization during natural viral infection. To ensure that the observed effect was not an artifact, cells were infected with ZIKV, and eIF4G1 levels were measured. Because it remains disputable whether a specific decrease in the level of a particular protein is reflected by changes in signal transduction, we tested the effect of ZIKV infection on the production of cellular proteins using the puromycin assay, with appropriate controls (32),(46). We did not observe any changes in host gene translation, proving that the effect on eIF4G1 cleavage is not likely to alter the physiology of the cell. However, modulation may occur locally at the replication site, and while it would improve viral replication, the effect on the whole cell may be too subtle to be detected. For this reason, the effect of eIF4G1 on ZIKV replication was tested by gene silencing and gene overexpression experiments, but the role of the eIF4G1 protein in viral replication remained elusive.

As listed in **Table 1**, several proteins have been reported as targets for the ZIKV NS3 protease. Intrigued by the results obtained for the eIF4G1 protein, we explored whether autophagy-related protein 16-1 (ATG16L1), c-Jun amino-terminal kinase-interacting protein 4 (JIP4), mitogen-activated protein kinase kinase kinase 7 (TAK1 or MAP3K7), disulfide-isomerase A3 (PDIA3), heterogeneous nuclear ribonucleoprotein A2/B1 (hnRNP A2/B1), aldolase A (ALDOA) (42), ER-localized reticulophagy receptor FAM134B (40), and septin-2 protein (41) may serve as targets for the NS3 protease. To our surprise, we could not confirm these previous observations, and we considered why this might be the case. First, in these previous studies, different expression systems and constructs were employed. In some cases, part of the NS2B cofactor was covalently linked to NS3 by a flexible linker. This is relevant, as it has been shown by others that the linker itself may alter the dynamics of the protein and, consequently, the substrate specificity of the protease. Second, the soluble version of the protease is not anchored at the membrane, which may also alter the substrate specificity. Third, the localization of the protease in the ER may limit the number of possible targets, and even proteins that may serve as NS3 substrates in biochemical assays may not be cleaved due to the differential spatial distribution. Finally, although mass spectrometry is sensitive enough to detect even minor changes in protein content, it could in some cases deliver results which are not relevant for the homeostasis of the intracellular environment. Therefore proteome changes detected by MS should be confirmed by other methods, and their biological relevance should be explored, using other approaches.

In conclusion, our study shows that the biological role of the ZIKV NS3 protease may be limited. This may be due to the relatively recent transmission of the virus on a large scale to the human population. It would therefore be interesting to analyze virus evolution in humans in the future, especially changes in the localization and/or substrate specificity of NS3 protease.

